# scLink: Inferring Sparse Gene Co-expression Networks from Single-cell Expression Data

**DOI:** 10.1101/2020.09.19.304956

**Authors:** Wei Vivian Li, Yanzeng Li

## Abstract

A system-level understanding of the regulation and coordination mechanisms of gene expression is essential to understanding the complexity of biological processes in health and disease. With the rapid development of single-cell RNA sequencing technologies, it is now possible to investigate gene interactions in a cell-type-specific manner. Here we propose the scLink method, which uses statistical network modeling to understand the co-expression relationships among genes and to construct sparse gene co-expression networks from single-cell gene expression data. We use both simulation and real data studies to demonstrate the advantages of scLink and its ability to improve single-cell gene network analysis. The source code used in this article is available at https://github.com/Vivianstats/scLink.

## Introduction

Biological systems often involve tens of thousands of genes tightly regulated in complex and dynamic networks, which could change substantially in different tissue types, developmental stages, or cell states [1, 2]. Therefore, elucidating gene interactions in a network manner is crucial for understanding complex biological processes in human physiology and pathology. By identifying abnormal gene interactions and regulations in disease states, it is possible to reveal the biological and biochemical pathways relevant to disease mechanisms and therapeutic targets [3]. For instance, transcriptional dysregulation revealed by disease-associated gene interactions has been reported in various diseases, including cancer [4, 5], neurological disorders [6], and psychiatric disorders [7], leading to functional insights of transcriptome organization in disease processes.

In network analysis, genes are represented by nodes, and their relationships are depicted by different types of directed or undirected edges between the nodes. The gene networks constructed from bulk tissue RNA sequencing (RNA-seq) data have played a key role in identifying genes that are responsible for similar biological functions, targets of transcriptional regulation, and regulators of disease-associated biological pathways [8, 9, 10]. However, the tissue-level networks can only describe the average gene-gene relationships across multiple biological samples [11]. Rapid advances of single-cell RNA sequencing (scRNA-seq) technologies have now made it possible to investigate gene networks across individual cells in a cell-type-specific manner [12]. ScRNA-seq technologies can parallelly profile gene expression levels in large numbers of individual cells, offering a unique opportunity for investigating genes’ relationships at a single-cell resolution. Based on the functional networks constructed from scRNA-seq data, biological discoveries have been made to provide novel insights into the transcriptional regulation mechanisms underlying various biological processes, including cancer progression [13], immune system response [14], and embryonic development [15].

Even though exploratory analyses demonstrated the possibilities of constructing functional gene networks across single cells, both technical and biological complications present challenges to the genome-wide inference of gene dependencies from scRNA-seq data [16]. Due to technical molecular inefficiencies, a truly expressed gene may not be detected by scRNA-seq in some cells, and thus is represented by a false zero expression level [17]. Meanwhile, the stochastic gene expression process can also lead to zero expression representing biological variation. Therefore, scRNA-seq data are often much sparser than the traditional bulk RNA-seq data, requiring new statistical and computational tools that could tackle the modeling challenges given the excess zero counts. In bulk RNA-seq data analysis, studies of gene networks mostly rely on the Pearson or Spearman’s correlation coefficients to characterize the gene co-expression strength [18, 19]. However, these two measures cannot provide a robust estimation of gene co-expression given the sparse scRNA-seq data with substantial technical noises and biological heterogeneity [20, 21].

In light of the aforementioned problem, Iacono et al. [19] used the correlation between two genes’ patterns of differential expression between cell types instead of gene expression levels to study gene regulatory network plasticity. However, most reconstruction methods of gene networks do not explicitly account for the sparsity issue. For example, PIDC uses partial information decomposition from the multivariate information theory to quantify the statistical dependencies between genes and infer gene networks from scRNA-seq data [22]. GENIE3 decomposes the prediction of a gene network between *p* genes into *p* different regression problems, and uses tree-based ensemble methods to infer the edges between genes [23]. It was shown to have competitive performance on bulk data [24] and has also been applied to single-cell data for gene network inference [25]. In addition to methods that are purely based on gene expression data, there are also single-cell methods developed to infer direct gene regulatory relationships instead of statistical dependencies [26]. To infer the direct gene interactions, these methods typically require external information such as time points or pseudo-time order of the cells [27, 28] and known transcription factors [25].

Despite being an active research area, accurate inference of functional gene networks from single-cell gene expression data remains a challenge, partly due to a lack of sufficient resolution in gene expression for making reliable inference [16, 26]. In this work, we propose a new method, scLink, to better characterize the statistical dependencies between genes in single cells, and improve the construction of gene co-expression networks based on a newly proposed co-expression measure. In summary, scLink has the following advantages. First, it proposes a robust estimator for measuring gene co-expression strength, built upon our previous work on improving the quality of single-cell gene expression data [29]. Instead of using all the observed read counts for measuring the association between two genes, scLink aims to rely on the cells in which both genes are accurately measured with high confidence. Second, scLink adapts the Gaussian graphical model [30] to distinguish the direct associations between genes from indirect ones and leads to easily interpretable sparse networks. Under this framework, the absence of an edge between two genes indicates the independence of these two genes conditioned on all other genes. Gaussian graphical models have been widely used to infer biological networks from genomic data and have revealed cancer-type-specific gene interactions that potentially contribute to cancer development and progression [31, 32, 33]. Third, scLink uses a penalized approach to identify relatively sparse gene networks in a data-adaptive manner, adjusting the penalty strength on each edge based on the observed co-expression strength in single cells. This penalized approach is a modified version of the graphical lasso method [34] to improve the identification of edges using single-cell data. We show that by combining the above features, scLink could enable more robust quantification of gene co-expression relationships, more accurate construction of gene co-expression networks, and better identification of functional gene modules that could provide insights into celltype-specific transcriptional regulatory mechanisms and molecular pathways.

## Methods

### A robust estimator for measuring gene co-expression strength

Accurate and robust estimation of the strength of gene co-expression relationships is the key to reliable inference of gene co-expression networks. Since single-cell gene expression data contain excess zero counts and relatively inaccurate low counts due to both technical and biological variability [35, 29], the conventional Pearson or Spearman’s correlation coefficients are often not reliable for single-cell gene expression data, especially for genes whose expression values are highly sparse (Figure 1) [36, 20]. In our previous work, we proposed a statistical method, scImpute, to address the excess zeros in scRNA-seq data [29]. Based on scImpute’s idea to identify the highly likely outliers (i.e., gene expression values that are not accurately measured), we propose a robust estimator for gene co-expression strength, which helps scLink to improve the inference of sparse gene co-expression networks.

**Figure 1:**
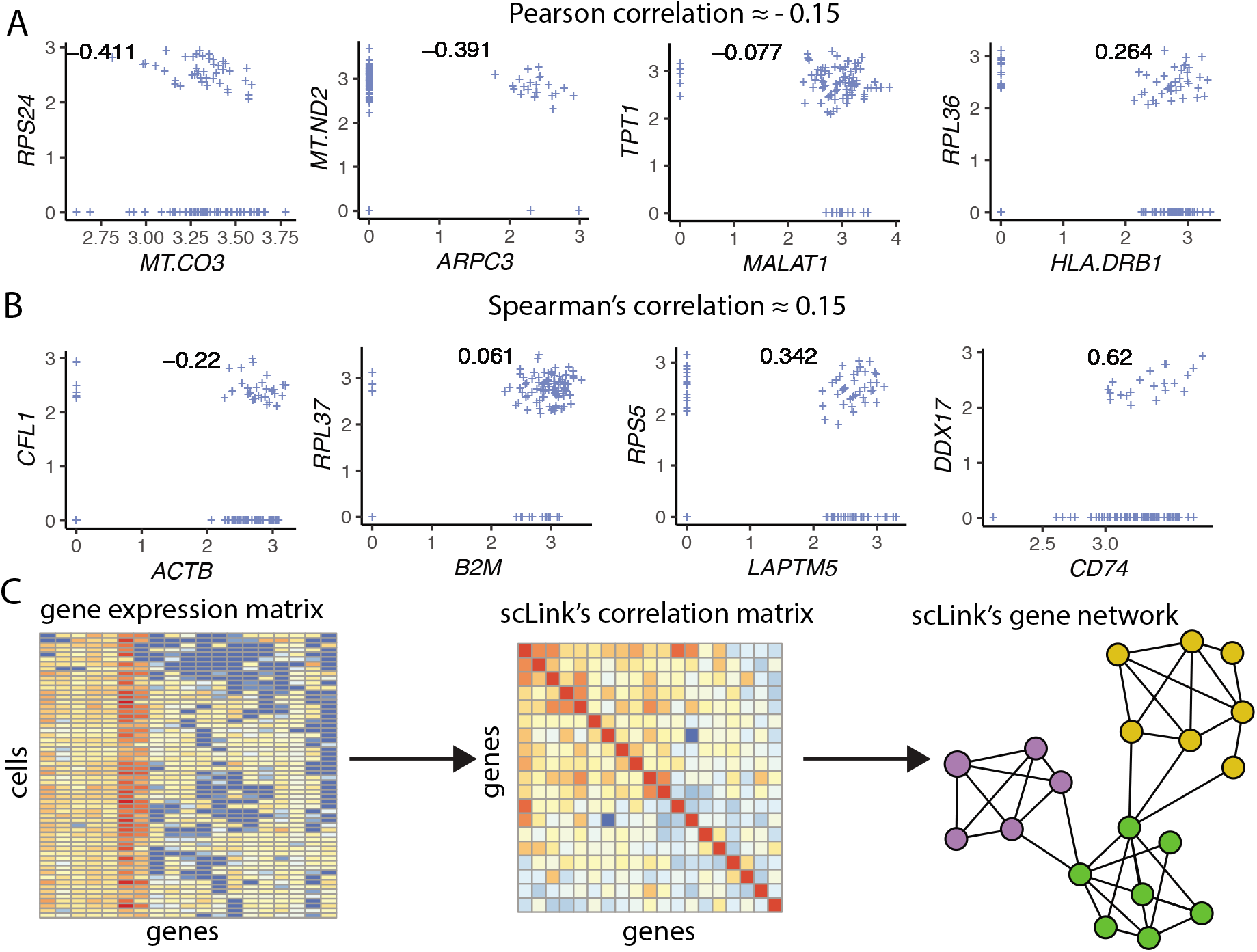
The motivation and workflow of the scLink method. **A-B**: Example gene pairs whose Pearson correlation (**A**) or Spearman’s correlation (**B**) are similar in the B cells. The scatter plots present the log_10_-transformed gene expression levels. The Pearson or Spearman’s correlation calculated using only cells in which both genes were detected are marked in the scatter plots. **C**: The workflow of the scLink method. In the first step, scLink calculates a robust coexpression matrix from the gene expression data. In the second step, scLink identifies a sparse gene network from the co-expression matrix using a penalized and data-adaptive likelihood approach.

Suppose the scRNA-seq data of a certain cell type (determined using biological markers or computational tools) is summarized as a read count matrix, with rows representing *n* cells and columns representing p genes. We normalize the count matrix by the library size of each cell, so that all cells have *M* reads after normalization. Typical choices for *M* include the median library size of all cells or a predetermined constant (e.g., 10^5^) [17]. Denoting the normalized matrix by ***C***, we apply the log_10_ transformation to the count matrix to prevent a few large observations from being extremely influential. The resulting matrix is denoted as ***Y***, with *Y_ij_* = log_10_(*C_ij_* + 1.01) (*i* = 1,2,…, *n,j* = 1,2,…, *p*). The pseudo count 0.01 is added to avoid infinite values in parameter estimation.

We also denote the log-transformed gene expression matrix without the pseudo-count as ***X***, where *X_ij_* = log_10_(*C_ij_* + 1) (*i* = 1,2,…, *n,j* = 1,2,…, *p*). In conventional methods, pairwise correlation coefficients are calculated from ***X*** to obtain the sample correlation matrix, based on which gene networks are constructed. In our scLink method, however, we add a filtering step to identify the accurately measured read counts and rely on these counts in network inference, by adapting a mixture model used in scImpute. Similar mixture models have been shown to effectively capture the bimodal characteristic of single-cell gene expression data [35, 37, 38]. Specifically, for each gene j, we assume its expression level is a random variable *Y_j_* following a Gamma-Normal mixture distribution, with a density function

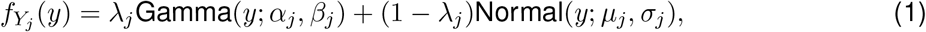

where λ_*j*_ is gene *j*’s non-detection rate, *α_j_* and *β_j_* are the shape and rate parameters in the Gamma distribution, and *μ_j_* and *σ_j_* are the mean and standard deviation in the Normal distribution. The Gamma distribution models the gene expression distribution when the sequencing experiments fail to accurately capture gene *j*’s transcripts, while the Normal distribution models the actual gene expression levels.

We designed an Expectation-Maximization algorithm to estimate the parameters in model (1), and these estimates are denoted as 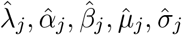, respectively. We can then filter the gene expression values based on the non-detection probability of gene *j* in cell *i*, which is estimated as

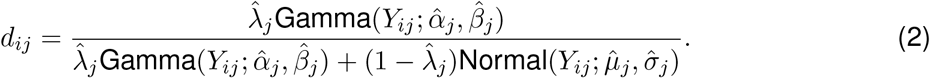

Since *d_ij_* ∈ (0,1) and a smaller *d_ij_* indicates better confidence of the observed gene expression *Y_ij_*, we can filter expression values by selecting a threshold *t*. Gene expression values whose corresponding d_ij_ < *t* are considered to be accurately measured with high confidence, while expression values whose corresponding *d_ij_* ≥ *t* are treated as missing values. We set *t* = 0.5 in our analysis, as we have previously demonstrated that the selection of this threshold only impact a tiny proportion of genes [29].

Given the identified accurate expression values and missing values, our robust estimator for measuring gene co-expression strength is defined as the pairwise-complete Pearson correlation coefficients. We calculate the co-expression strength from gene expression matrix ***X***, where *X_ij_* = log_10_(*C_ij_* + 1) (*i* = 1,2,…, *n,j* = 1,2,…, *p*). For genes *j*_1_ and *j*_2_, their robust correlation is calculated as

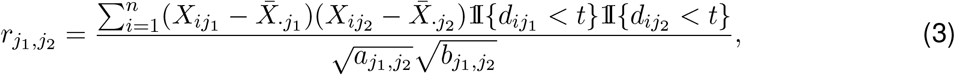

where 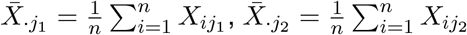, and

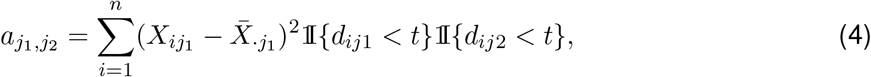

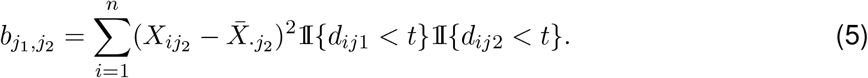

The pairwise robust correlation coefficients are used by scLink to construct gene co-expression networks in the following subsection. To improve the robustness of our analysis, if the sample size for calculation between genes *j*_1_ and 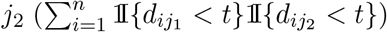 is smaller than 10, then we instead use the Pearson correlation coefficient for this pair of genes.

### The scLink method for gene network inference

To construct sparse gene co-expression networks from single-cell gene expression data, our scLink method adapts the Gaussian graphical model [30] and the penalized likelihood method [39, 34], which uses the principle of parsimony to select the simplest graphical model that adequately explains the expression data. We assume that the actual gene expression values in each cell, without missing values being present due to technology limitations, be a *p*-dimensional random vector ***Z*** = (*Z*_1_,…, *Z_p_*)^*T*^ following a multivariate distribution *N* (***μ***, **Σ**). Note that ***Z*** denotes the actual gene expression on the log_10_ scale, and ***Z*** is a hidden variable that is not directly observable. We wish to estimate the concentration matrix **Θ** = **Σ**^−1^, since a zero entry *θ*_*j*1;*j*2_ = 0 indicates the conditional independence between the two genes *j*_1_ and *j*_2_ given all other genes. In other words, if we consider an undirected graph *G* = (*V,E*), where *V* contains p vertices corresponding to the *p* genes and the edges are denoted by *E* = {*e*_*j*1;*j*2_}_1≤*j*1≤*j*2≤*p*_, then the edge between genes *j*_1_ and *j*_2_ is absent if and only if *θ*_*j*1;*j*2_ = 0. Thus we can infer the presence of edges between genes by estimating parameters and identifying non-zero entries in the concentration matrix **Θ**.

Given a random sample (*n* cells) of ***Z***, a commonly used lasso-type estimator [34] takes the form

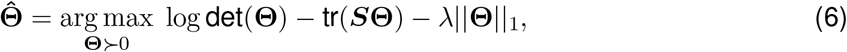

where (logdet(**Θ**) – tr(**SΘ**)) is proportional to the log-likelihood for **Θ** (ignoring a constant not depending on **Θ**) [39] and 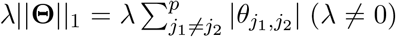 is a penalty term adding a constraint on the number of non-zero elements in the concentration matrix. In model (6), ***S*** denotes the estimated covariance matrix.

Recall that we summarize the observed gene expression matrix as **X**, where *X_ij_* = log_10_(*C_ij_* + 1) (*i* = 1,2,…, *n,j* = 1,2,…, *p*). If we directly consider each matrix column **X**._1_…, **X**._*j*_ as a realization of ***Z***, the covariance matrix could be estimated with the sample covariance matrix 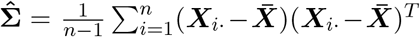, where 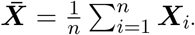. However, due to limited detection capacity in scRNA-seq technologies as we have discussed in the previous subsection, the observed gene expression values ***X***._1_,…, ***X***._*j*_ cannot be directly treated as a sample of ***Z***, and 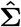 is not an ideal estimator of the covariance matrix **Σ**. In scLink, we estimate **Σ** with a robust estimator **S**, which is constructed with the robust estimator for gene co-expression strength as introduced in the previous subsection. The elements in ***S*** are calculated as

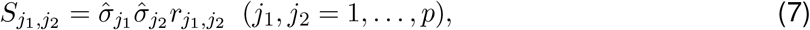

where *r*_*j*1;*j*2_ is the robust correlation given by formula (3), and 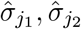 are respectively the estimated standard deviation of genes *j*_1_ and *j*_2_ from the mixture model (1). This idea of robust covariance estimation is motivated by a general framework for robust covariance calculation of high-dimensional data [36], and implemented with careful consideration of single-cell data characteristics.

In addition to proposing the robust estimator of covariance matrix, another improvement by scLink is to introduce a data-adaptive penalty term instead of using a constant λ. We expect the penalty to be stronger on *θ*_*j*1;*j*2_ if the robust correlation between genes *j*_1_ and *j*_2_ is weaker, and vice versa. Therefore, we propose a weighted penalty term 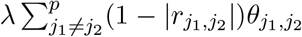 to incorporate gene-pair-specific information when adding the sparsity constraint on estimated concentration matrix 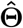.

In summary, the scLink estimator of the concentration matrix takes the form

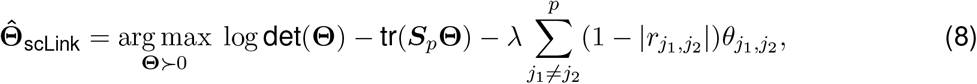

where λ > 0 and ***S***_*p*_ is a positive semidefinite approximation of ***S***, which is defined in (7). In detail, ***S***_*p*_ = ***S*** + | min{0,*τ*}***I***, where *τ* is the smallest eigen value of ***S*** and ***I*** is the identity matrix [36]. There are multiple algorithms that can be implemented to solve model (8), and we selected the QUIC algorithm [40] since its computational cost is *O*(*p*) and has a superlinear convergence rate. After we obtain the estimated 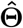, for network construction it follows that 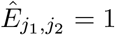 if 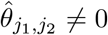 and 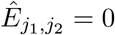 if 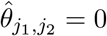.

### Selection of the regularization parameter

In model (8), the value of the regularization parameter λ would influence the level of sparsity in the estimated concentration matrix and therefore the constructed gene network. Here we discuss two approaches that can be used to guide the selection of λ. The first approach is to use the Bayesian information criterion (BIC) [41]. For model (8) and a particular value of λ, BIC is calculated as

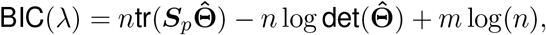

where *m* is the total number of edges (i.e., non-zero elements in 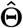). We can apply model (8) on single-cell gene expression data with a sequence of varying regularization parameters, and select the value of λ that leads to the minimal BIC value. The second approach is to directly select λ based on the level of sparsity in the constructed gene networks. Suppose we have an expectation for the sparsity level based on prior knowledge (e.g., existing biological networks or pilot studies), we can select the value of λ that achieves the expected sparsity level of the gene network.

### Simulation of synthetic gene networks and expression data

We adapted the procedures described in Mestres et al. [42] to simulate network structures. In each simulation setting, we first generated a block diagonal connectivity matrix ***E***_*p*×*p*_, where each block had a hub-based or power-law topology and the whole matrix also contained a fixed number of random connections between blocks. In the connectivity matrix, |*E*_*j*1;*j*2_| = 1 means that there’s an edge between genes *j***1** and *j***2**, and *E*_*j*1,*j*2_ = 0 means that there’s no edge between the two genes. This process was assisted with the R package `ldstatsHD v1.0.1` [42]. Given the connectivity matrix, a partial correlation matrix was simulated by

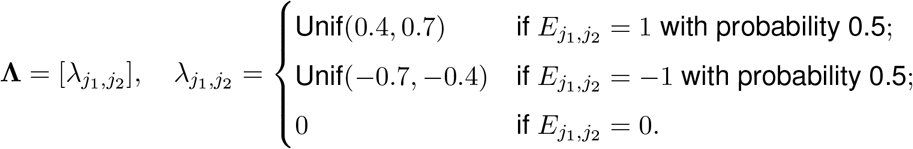

In case that **Λ** was not positive definite, we applied the transformation **Λ**_*p*_ = **Λ** + | min{0,*τ*}|***I***, where *τ* was the smallest eigen value of **Λ** and ***I*** was the identity matrix. We then calculated the corresponding correlation matrix ***R***, and used it together with the gene expression mean and standard deviation estimated from a real scRNA-seq dataset [43] to simulate synthetic gene expression matrix ***X***^0^ from a Multivariate Gaussian distribution. The estimation of the gene expression parameters followed the procedures described in the Methods section: A robust estimator for measuring gene co-expression strength. Next, we introduced zero counts to the gene expression matrix to mimic the observed zeros due to technical missing values and biological non-expression. Since the possibility of observing zero counts for a gene is negatively correlated with this gene’s mean expression in real data [35, 29], we calculated a probability for each entry in the gene expression matrix: 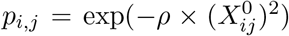, where *ρ* was a parameter controlling the dependence between missing probability and gene expression. Then a binary indicator was sampled for each entry: *I_i,j_* ~ Bernoulli(*p_i,j_*), with *I_i,j_* = 1 indicating that the corresponding entry would be replaced by 0. In other words, we assumed *p_i,j_* was the probability of observing a zero count for gene *j* in cell *i*. Therefore, the final single-cell gene expression matrix was defined as ***X***, where 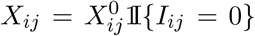. Repeating the procedures described above with different values of *ρ*, we could generate synthetic single-cell gene expression matrices with known network topologies and different levels of sparsity. In our study, we used four different values of *ρ*: {0.07,0.10,0.13,0.16}.

### Calculation of the robustness score

We use scLink as an example to describe how the robustness score was calculated in the Results section. The robustness of PIDC and GENIE3 were calculated using the same method. Suppose for a specific cell number (*n*) and gene number (*p*), by randomly sampling n cells from the given cell type for *L* times (*L* = 10 in our analysis), we obtained *L* gene adjacency networks from scLink: *E*^(*ℓ*)^ (*ℓ* ∈ {1,2,…, *L*}), where 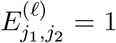 if the two genes *j*_1_ and *j*_2_ have an edge in the *ℓ*-th gene co-expression network; otherwise, 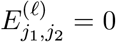. To simplify the notation, we denote 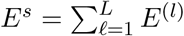. The robustness of scLink is then calculated as

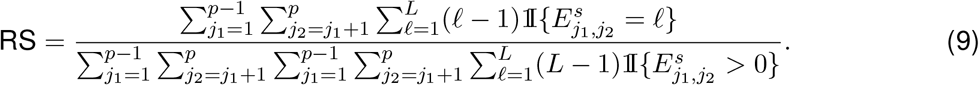

For example, RS = 1 if the *L* inferred adjacency networks are exactly the same; RS = 0 if the *L* gene networks do not have any overlap.

## Results

### The scLink method

To improve the construction of gene co-expression networks for single cells, we propose the scLink method to calculate the correlation between gene pairs, and then use a penalized and data-adaptive likelihood method to learn sparse dependencies between genes and construct sparse gene co-expression networks. One motivation of scLink is that the conventional Pearson and Spearman’s correlation coefficients do not provide an efficient approach to representing and interpreting gene associations given the high sparsity of single-cell gene expression data. For instance, we calculated the Pearson and Spearman’s correlation for a gene expression dataset of 109 immune B cells [13] (Supplementary Figure S1), after normalization and log transformation as described in Methods. We only used the 410 genes with at least 10% detection rates, and the proportion of zero counts in the expression matrix is 71.2%. As an example, there are 886 gene pairs with a similar Pearson correlation coefficient in [−0.15, −0.14], and 1302 gene pairs with a similar Spearman’s correlation in [0.14,0.15], but their association varies in a much larger range when we recalculated the correlation only using the cells in which both genes were detected (Figure 1A-B). To more accurately infer gene co-expression networks that can capture functional gene modules, scLink has two key steps (Figure 1C, Methods). The first step is to calculate a robust co-expression matrix from the gene expression data to accurately represent the co-expression relationships among the genes. In this step, scLink proposes a robust estimator for measuring gene co-expression strength, while relying on the cells in which both genes are accurately measured with high confidence. In the second step, scLink aims to identify a sparse gene network from the co-expression matrix using a penalized and data-adaptive likelihood approach. We use both simulation and real data studies to demonstrate the efficiency of scLink, and how scLink improves the identification of functional gene modules, regulatory relationships, and co-expression clues about cellular mechanisms that are active in normal and disease processes.

### scLink demonstrates efficiency in simulation studies

Our motivations for using simulated scRNA-seq data based on synthetic networks are two-fold. First, since the actual gene networks underlying real single-cell gene expression data are unknown, the synthetic networks provide ground truth for comparing computational methods in a systematic and unbiased manner. Second, using the simulated data, we can evaluate the performance of gene network inference methods given diverse network architectures and experimental settings. These results can help us investigate the advantages of each method in different scenarios.

In our simulation setting, we considered two types of network topology: hub-based networks and power-law networks [42]. The power-law networks (Figure 2A) assume that the distribution of the node degrees (i.e., the total number of edges a node has) follows a power law [44]. That is, *a*(*k*) = *k*^*−α*^/*ζ*(*α*), where *a*(*k*) denotes the fraction of nodes with degree *k, α* is a positive constant, and *ζ*(·) is the Riemann zeta function. In contrast, in the hub-based networks (Figure 2B), a few nodes have a much higher degree than the rest nodes, and these high-degree nodes represent hub genes with critical functions in biological networks [18]. Using a carefully designed simulation framework (see Methods), we could generate synthetic single-cell gene expression matrices with known network topologies and different levels of sparsity. Therefore, we can evaluate the accuracy of a computationally inferred gene network by comparing it with the ground truth network.

**Figure 2:**
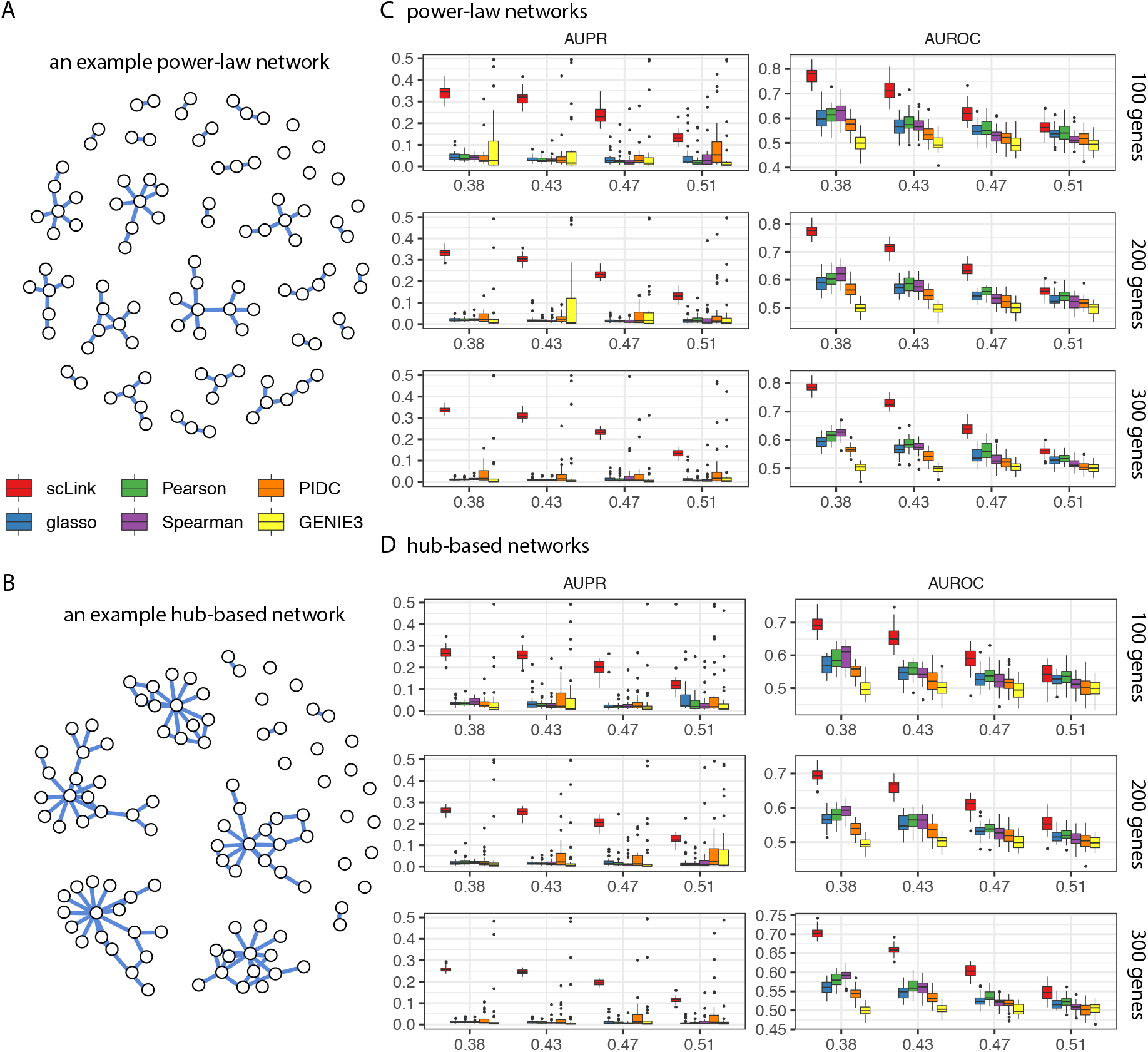
Comparison of scLink and the other gene network inference methods on synthetic single-cell gene expression data. **A**: An example 100-gene network with the power-law topology. **B**: An example 100-gene network with the hub-based topology. **C**-**D**: AUPR and AUROC scores of scLink and the other five methods given gene expression data generated from the power-law networks (**C**) or hub-based networks (**D**). The gene expression matrices have varying number of genes (100,200, or 300) and proportion of zero counts (marked on the *x*-axis).

We compared scLink with five alternative methods on the synthetic data to evaluate their accuracy in constructing gene co-expression networks. Among these five methods, PIDC infers genes’ dependencies in single cells based on the multivariate information theory [22]. GENIE3 was first developed to infer regulatory networks from bulk expression data [23], and was recently applied to single-cell expression data [25]. These two methods were demonstrated to have leading performance on simulated and real scRNA-seq data in a recent comparison [16]. In addition, our comparison also included glasso, a general statistical method for estimating sparse networks [34], and its variants have been used to address different challenges in gene network construction [33, 45]. Finally, we also considered gene networks constructed by thresholding the Pearson or Spearman’s correlation coefficients, as these are commonly used statistical measures for constructing gene co-expression networks [18].

Since the gene networks are expected to be sparse, we use the area under the precision-recall curves (AUPR) as the primary criterion and the area under the receiver operating characteristic curves (AUROC) as the secondary criterion, to achieve a fair and comprehensive comparison of the methods. For scLink and glasso, the curves were obtained by evaluating the two methods with different values of the regularization parameter (i.e., λ, see Methods). For PIDC and GENIE3, the curves were obtained by thresholding the estimated edge weights (between every pair of genes) at different values. For Pearson and Spearman’s correlation, the curves were obtained by thresholding the absolute values of correlation coefficients. For each type of network topology, we simulated synthetic single-cell gene expression data with 100, 200, and 300 genes (nodes). By changing the simulation parameters, we could generate gene expression data with different sparsity level (i.e., the proportion of zero counts). In each parameter setting, the simulation was independently repeated 50 times. We compared the computationally inferred gene networks with the true underlying gene networks using the AUPR and AUROC scores.

Our comparison results show that for both power-law and hub-based network topologies, scLink has the best AUPR and AUROC scores, outperforming glasso, PIDC, GENIE3, and the two correlation measures (Figure 2C-D). We think an important reason why scLink demonstrates higher accuracy is that it explicitly models the excess zero or low counts in single-cell data, providing a more robust estimator for gene co-expression strength to be used in the network inference step. Evaluating the performance of glasso, we found that it has a similar or even slightly lower accuracy compared with directly thresholding the Pearson correlation coefficients. This implies that the basic penalized Gaussian graphical model is not very efficient for single-cell data. However, by incorporating improved co-expression measures and adding data-adaptive penalties to the genegene edges, scLink largely improves the accuracy of the graphical model. We also observed that most methods, including scLink, have increasing accuracy on less sparse single-cell gene expression data. A primary reason is that these datasets provide a larger effective sample size for network inference and contain fewer noises that could lead to false discoveries. This result suggests that it could be advantageous to filter out lowly expressed genes before network inference for real single-cell gene expression data. In addition, given the same level of sparsity in single-cell data, most methods tend to have better performance on the power-law networks than the hub-based networks. A possible reason is that when multiple genes are simultaneously interacting with the same hub gene, it is very challenging to precisely distinguish the direct dependencies from the indirect ones among these genes only using the gene expression data.

As a proof-of-concept study, we also compared two modified versions of glasso with scLink on the simulated data. The first method, glasso-r, refers to the glasso method based on scLink’s robust correlation measure. It is the same as scLink except that it uses a uniform weight of 1 instead of the adaptive weights in the penalty term. In other words, the penalty term is replaced with 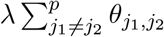 in model (8). The second method, glasso-f, refers to the glasso method with a filtering procedure. It filters out the cells that have greater than *x*% of zero counts before applying the glasso approach, and we set *x* = 70 in this study. This reflects the practice to filter out low-quality cells in real data applications. Our results based on both power-law and hub-based networks show that glasso-f does not effectively improve the network construction accuracy compared with glasso (Supplementary Figure S2). In addition, scLink achieves higher AUPR than glasso-r while the two methods have similar AUROC, suggesting the additional benefit of using adaptive penalty for constructing single-cell gene networks. Furthermore, as a control study, we also compared scLink with the other methods on simulated data without introducing an extra level of sparsity (Supplementary Figure S3). In this control study, the synthetic scRNA-seq data were also generated as described in Methods, except that the step of introducing zero counts is skipped. As expected, all the methods, especially the two methods based on Pearson and Spearman’s correlation, have improved accuracy compared with the performance on sparse gene expression data. The control study demonstrates the unique challenge presented by the high level of sparsity in single-cell gene expression data, and the need to develop specific methods accounting for these data characteristics in the modeling step.

### scLink identifies cell-type-specific gene networks from the Tabula Muris data

To evaluate scLink’s performance on experimental single-cell data and demonstrate its application to construct cell-type-specific gene networks, we applied scLink to gene expression values derived from Smart-Seq2 RNA-seq libraries [46]. This dataset from the Tabula Muris database includes 53,760 cells of 20 different tissues from 8 mice [47], providing a valuable opportunity to perform the analysis in a cell-type-specific manner. We separately applied scLink to gene expression datasets of 59 cell types (each with at least 100 cells), using the top 500 highly expressed genes in each dataset. The proportion of zero counts in the 59 gene expression matrices ranges from 1.0% to 40.5%, and has a mean of 12.4%. The regularization parameters (λ) in model (8) were selected as the smallest value from {1,0.95,…, 0.05} such that the resulting network had no more than 5% edges (i.e., 5% × (500 × 499)/2 ≈ 6237 edges). After applying scLink to construct the cell-type-specific gene co-expression networks, we summarized the gene degrees, number of network communities (identified by the Louvain algorithm [48]), and sizes of the communities in Supplementary Figure S4.

Since the true underlying gene networks are unknown for real gene expression data, we first investigated the identified edges between genes and known transcription factors (TFs). For each cell type, we calculated the number of identified edges connected to known TFs in scLink’s results, and assessed their overlap with the TF-target edges discovered in previous ChIP-seq experiments [49, 50, 51]. Since the ChIP-seq experiments were performed using bulk data from human or mouse tissues instead of single cells, we pooled the TF-target pairs from the ChIP-seq experiments for the comparison, resulting in a database of 310 TFs. We used this database as a reference to investigate scLink’s performance, but we note that it’s not appropriate to calculate precision and recall rates treating this database as the ground truth. Our results show that a substantial proportion of the identified TF-gene edges by scLink were previously discovered in ChIP-seq experiments (Figure 3A). This proportion ranges from 15.6% to 89.5% among different cell types and has a median of 59.3%. Especially, the scLink identification has relatively high consistency with the ChIP-seq database in the epithelial, mesenchymal, pancreatic, epidermal, and muscle cell types, with a median overlapping proportion of 65.5%, 69.8%, 64.1%, 61.0% and 65.4%, respectively (Figure 3A). As a comparison, we also applied five additional network construction methods described in previous simulation studies to the Tabula Muris data: Pearson correlation, Spearman’s correlation, PIDC, glasso-f, and glasso-r (Supplementary Table S1). The median proportions of identified TF-gene edges that were previously discovered in ChIP-seq experiments are 53.0%, 53.6%, 63.5%, 56.4%, and 58.1% for these methods, respectively. We found that scLink and PIDC generally lead to a higher consistency with the bulk tissue ChIP-seq database.

**Figure 3:**
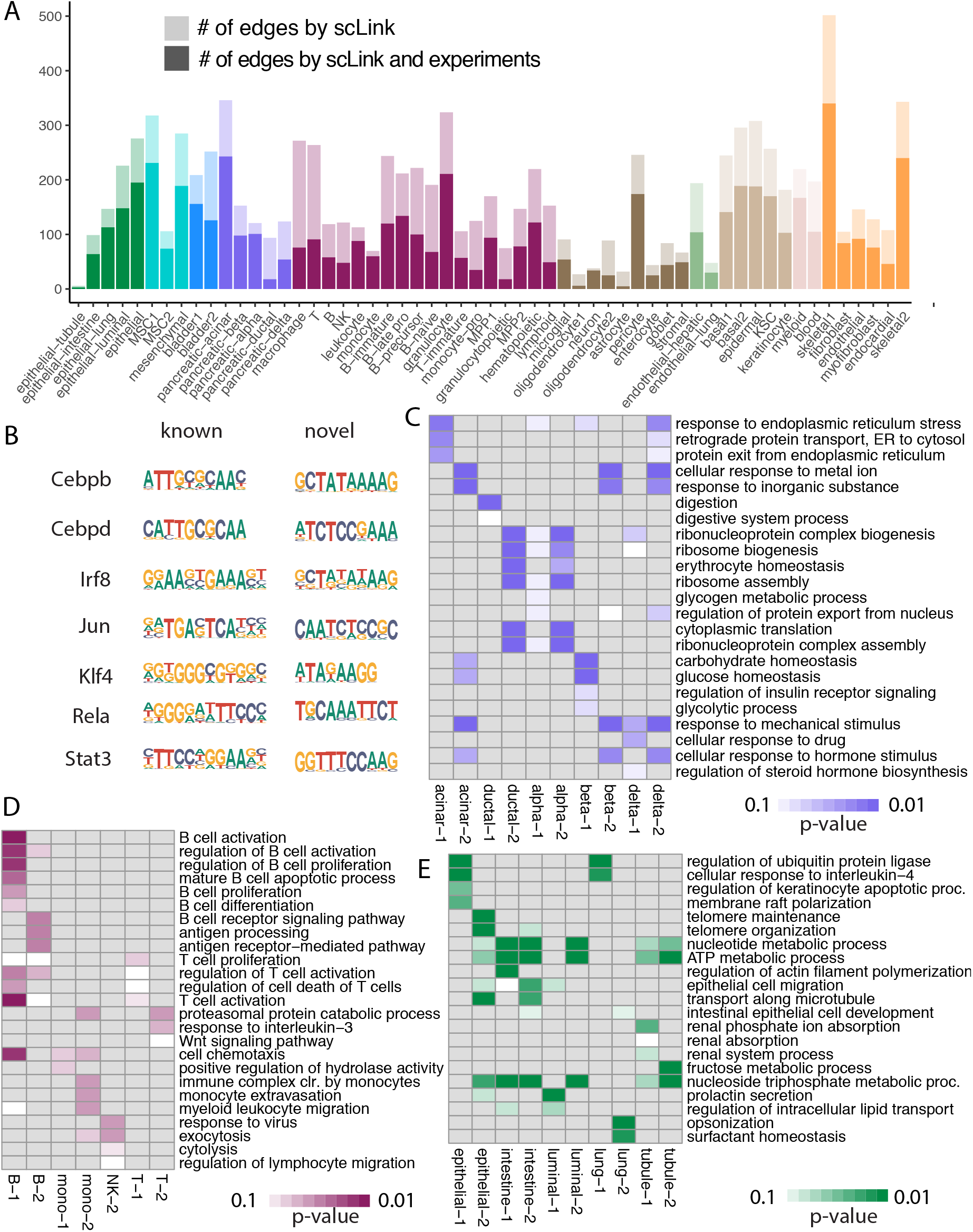
The performance of scLink on the Tabula Muris dataset. **A**: The numbers of TF-gene edges identified by scLink only and by both scLink and ChIP-seq experiments. **B**: Known and novel motifs of seven TFs identified from the promoter regions of genes connected to these TFs by scLink. **C-E**: Enriched GO terms in the two largest gene modules identified from pancreatic cell types (**C**), immune cell types (**D**), and epithelial cell types (**E**). FDR-adjusted *p* values are shown in the heatmaps.

Since some TF-gene edges identified by scLink were not previously observed from ChIP-seq experiments, we performed an additional motif analysis using HOMER [52] to study if the genes connected to the same TF by scLink share common motifs in their promoter regions. Our motif analysis identified both known and novel motifs for a group of TFs, including *Cebpb, Cebpd, Irf8, Jun, Klf4, Rela,* and *Stat3* (Figure 3B). Although we cannot conclude that these genes connected to the TFs by scLink are the direct targets of these TFs, the motif analysis shows that the genes connected to the same TF do have shared sequence features in their regulatory sequences and are likely to be co-expressed. In summary, the above analyses demonstrate that by estimating the single-cell gene networks, scLink is able to identify edges between TFs and their potential target genes, even though scLink does not rely on any prior information of known TFs.

To further validate the biological functions of gene co-expression networks estimated by scLink, we investigated the gene modules present in the gene networks for the pancreatic, immune, and epithelial cell types. These modules are supposed to represent groups of highly co-expressed genes that share similar biological functions or pathways in the corresponding cell types. For each cell type, we calculated the partial correlation matrix for the genes based on the estimated concentration matrix 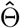 by scLink (see Methods). Then, we performed hierarchical clustering using (1-|partial correlation|) as the distance measure. Next, the genes were divided into separate modules by cutting the dendrogram at the height of 0.85. Finally, we performed the Gene Ontology (GO) enrichment analysis on the gene modules.

The enriched GO terms in the two largest gene modules of each cell type are displayed in Figure 3C-E. For the pancreatic cells (Figure 3C), we found that GO terms related to protein transportation and digestive system process are enriched in gene modules of exocrine cells (acinar and ductal cells), while GO terms related to glycogen metabolic process, glucose homeostasis, and cellular response to hormone stimulus are enriched in gene modules of endocrine cells (alpha, beta, and delta cells). It is worth noting that “regulation of insulin receptor signaling pathway” is only enriched in a gene module of beta cells, which have a critical role in insulin regulation [53]. In contrast, “regulation of steroid hormone biosynthetic process” is only enriched in a gene module of delta cells, which secrete the hormone somatostatin [54]. For the immune cells (Figure 3D), we found that GO terms related to B cell activation or proliferation are enriched in the largest gene module of B cells, and the terms related to B cell receptor signaling pathway and antigen processing are enriched in the second-largest module of B cells. In contrast, GO terms enriched in monocyte gene modules are related to monocyte extravasation and monocyte immune complex clearance. For the epithelial cells (Figure 3E), the enriched GO terms in gene modules also demonstrate cell-type-specific biological functions. For example, “surfactant homeostasis” is enriched in a module of lung epithelial cells [55], “prolactin secretion” is enriched in a module of luminal epithelial cells (of the mammary gland), and “intestinal epithelial cell development” is enriched in a module of large intestine epithelial cells.

To investigate if the gene-gene edges identified by scLink could improve the identification of molecular pathways and functional gene modules, we studied the inferred gene networks in T cells, skeletal muscle satellite stem cells, and pancreatic beta cells as three examples. In each inferred gene network by scLink, we focused on the largest connected component supported by known protein interactions in the STRING database [56], and we found that scLink can identify gene interactions that would be missed using a conventional approach with the Pearson or Spearman’s correlation (Supplementary Methods). In T cells (Figure 4A), the inferred network by scLink contains four modules corresponding to gene sets of different functions. However, these four modules are not reported in correlation-based networks with the same level of sparsity. The smallest module contains three genes associated with the GO term “protein serine/threonine phosphatase complex”, which was shown to be a requisite of T cells’ functions [57]. The second module contains three genes (*Rac1, Cdc42,* and *Cyba*) in the pathway of leukocyte transendothelial migration [58]. The two largest modules each contain nine genes, responsible for cell adhesion/receptor and antigen processing, respectively. Genes in the cell adhesion module are involved in the pathways of cell adhesion, cell surface interactions, and cell surface receptors; genes in the antigen processing module are responsible for antigen processing and presentation and immunoregulatory interactions [58]. In the muscle stem cells (Figure 4B), the inferred network contains a module of nine genes playing key roles in osteoclast differentiation [59, 60]. In this module, three edges (*Jund-Fos, Fosb-Fosl2, Fosl1-Fosl2*) are only identified by scLink. In addition, the network also contains a four-gene module involved in muscle regeneration [61, 62] and a five-gene module which was shown to play roles in myoblast differentiation [63, 64, 65]. These two modules are also missed by the classic correlation approach. In the pancreatic beta cells (Figure 4C), the inferred network identifies a module of 15 genes responsible for oxidative phosphorylation, which plays an important role in beta cells’ proliferation, survival, and response to rising blood glucose [66]. The above results demonstrate that scLink is able to identify edges between genes with direct interactions or close biological functions in a cell-type-specific manner. For an additional comparison, we also investigated the largest connected components supported by known protein interactions in the gene networks based on Pearson or Spearman’s correlation, PIDC, glasso-f, and glasso-r of the three cell types (Supplementary Methods). The largest gene modules in these networks are enriched with genes coding for ribosomal proteins (Supplementary Table S2) and/or are enriched with GO terms of biosynthetic and metabolic processes (Supplementary Tables S3-S7). These results show that the largest gene modules identified by these methods capture less cell-type-specific information and functional relevance than those gene modules identified by scLink.

**Figure 4:**
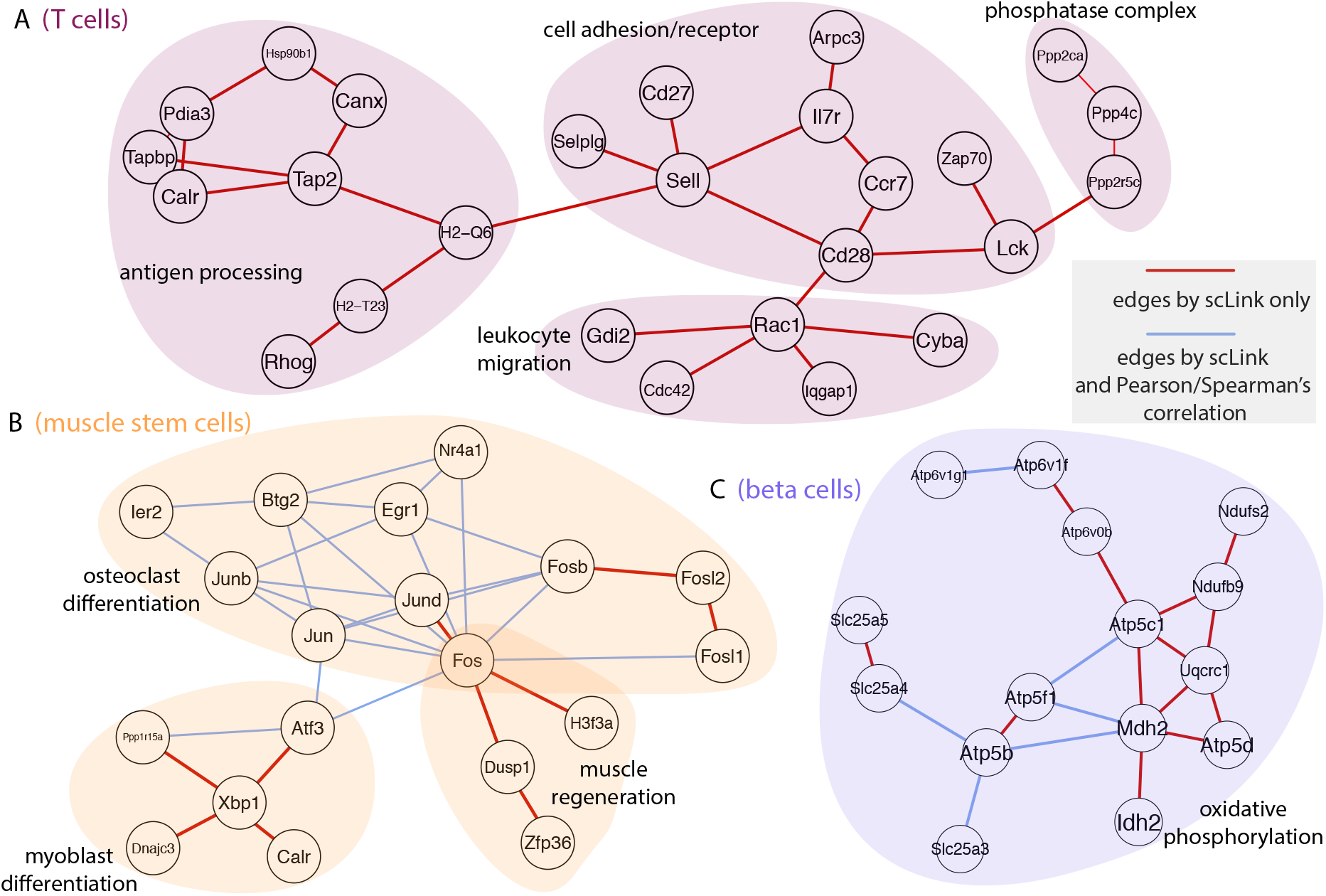
Gene networks inferred by scLink overlap with functional protein interaction networks. Inferred edges by scLink (red and blue) are displayed for T cells (**A**), skeletal muscle satellite stem cells (**B**), and pancreatic beta cells (**C**). Edges that are also identified by the correlation approach are displayed in blue. All the displayed edges are supported by known protein interactions in the STRING database. Functional gene modules are grouped in the shaded area.

### scLinks identifies gene network changes in breast cancer

We next applied scLink to a breast cancer single-cell dataset to study if scLink can assist with comparing gene networks in health and disease states. We collected the gene expression data of immune cells in the tumor (656 cells) and matched breast tissue (211 cells) from the same patient [13]. These expression values were measured by the inDrop platform [67]. We separately applied scLink to data from the normal and tumor tissue, using the top 500 highly expressed genes in the normal tissue. The proportions of zero counts in the normal and tumor expression matrices are 49.0% and 66.0%, respectively. The regularization parameters (λ) for scLink were selected as the smallest value from {1.2,1.1,…, 0.5} such that the inferred network has no more than 5% edges (i.e., 5% × (500 × 499)/2 ≈ 6237 edges). In the identified normal tissue gene network, the gene degree ranges from 2 to 103 with an average of 24.6, and the 89 communities identified by the Louvain algorithm has an average size of 5.6. In the tumor tissue gene network, the gene degree ranges from 2 to 137 with an average of 26.2, and the 63 communities identified by the Louvain algorithm has an average size of 7.9.

By comparing the inferred gene networks of the normal and tumor tissues, we found 453 differential edges (with a greater than 0.5 change in scLink’s correlation measures) that are only present in the normal cells but not in the tumor cells (Supplementary Figure S5A). We assessed the statistical significance of scLink correlation for the 453 edges in the normal condition using a bootstrap approach (Supplementary Methods), and 84.5% of the edges have a *p* value < 0.05 after adjusting for the false discovery rate (FDR) (Supplementary Figure S6). For example, in the normal cells, *FNBP1* is co-expressed with *MGP,* whose down-regulation is associated with better survival in breast cancer [68] (scLink’s correlation = 0.77, adjusted *p* = 0) (Figure 5A). However, they expressed in the tumor cells in a mutually exclusive manner (scLink’s correlation = −0.03). As another example, scLink identifies an edge between *EGR1* and *NUDT3* in normal cells (scLink’s correlation = −0.48, adjusted *p* = 0.004) but not in the tumor cells (scLink’s correlation = 0.35), and both genes were reported to have regulatory roles in breast cancer (Figure 5A) [69, 70].

**Figure 5:**
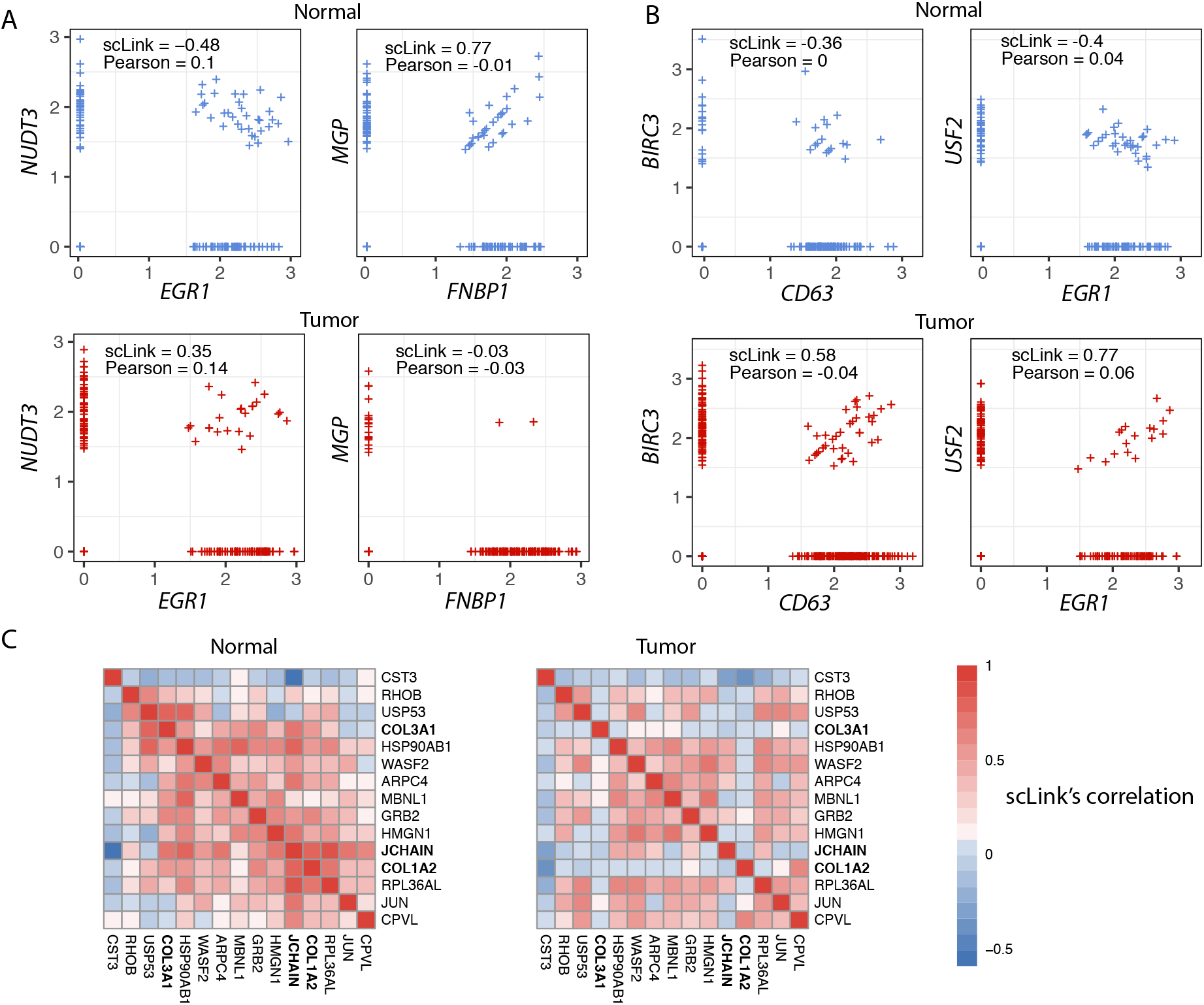
scLink identifies differential co-expression relationships in normal and breast cancer tissues. **A**: The expression of two gene pairs (*NUDT3* and *EGR1; FNBP1* and *MGP*) are highly associated in normal immune cells but not in breast cancer immune cells. Both scLink’s and Pearson correlation coefficients are displayed. **B**: The expression of two gene pairs (*CD63* and *BIRC3; EGR1* and *USF2*) are highly associated in breast cancer immune cells but not in normal immune cells. Both scLink’s and Pearson correlation coefficients are displayed. **C**: The correlation matrices of a 15-gene module identified from the normal immune cells by scLink.

Meanwhile, we identified 1384 differential edges (with a greater than 0.5 change in scLinks correlation measures) that are only present in the tumor cells but not the normal cells (Supplementary Figure S5B). We also assessed the statistical significance of the scLink correlation for the 1384 edges in the tumor condition, and 90.0% of the edges have a *p* value < 0.05 after adjusting for the FDR (Supplementary Figure S6). For instance, *EGR1* and *USF2* are highly co-expressed in tumor cells (scLink’s correlation = 0.77, adjusted *p* = 0.010) but not in normal cells (Figure 5B). A similar expression pattern was observed between *CD63* and *BIRC3* (scLink’s correlation = 0.58, adjusted *p* = 0 in tumor cells). In addition to *EGR1, CD63* and *BIRC3* were also found to be associated with breast cancer, while *USF2’s* role in breast cancer has not been clearly investigated [71, 72]. The above results demonstrate that by comparing co-expression changes between health and disease states, it is possible to (1) identify new genes that are associated with a particular type of disease; (2) investigate how co-expression and co-regulation of genes impact cell functions [13]. In contrast, the above co-expression changes between these gene pairs cannot be captured by the Pearson correlation coefficients, since their values remain low on the sparse single-cell expression data (Figure 5A-B). Actually, the Pearson correlation changes by no more than 0.2 for 97.0% of the edges between the normal and tumor states (Supplementary Figure S5C), implying that it is not sensitive for identifying important co-expression changes in single cells. For a more systematic comparison, we constructed Pearson correlation networks with the same level of sparsity (i.e., 6237 edges). We then investigated the biological functions of the 50 genes with the largest degree changes between the normal and breast cancer conditions, using the scLink and Pearson correlation networks. We found that the top enriched GO terms in the 50 genes identified by scLink are closely related to immune responses of myeloid cells, leukocytes, and neutrophils, while those enriched in the 50 genes identified by Pearson correlation are relevant to more general protein regulation processes and humoral immune response (Supplementary Table S8).

By comparing the tight gene modules embedded in the gene networks of normal and tumor cells, we could observe a dramatic change in the global network structure in addition to the change in individual edges. By performing hierarchical clustering using the partial correlation matrix estimated by scLink, we identified 12 modules with more than 10 genes in the immune cells of normal breast tissue (Supplemental Figure S7). Similarly, we identified 11 modules with more than 10 genes in the immune cells of tumor condition (Supplemental Figure S8). However, the two sets of module assignments only have an adjusted Rand index of 0.10 and a normalized mutual information of 0.49, implying widespread rewiring of gene networks in the breast cancer condition [73]. For example, a 15-gene module identified from the normal condition is much less densely connected in the tumor condition (Supplementary Figure S9). In this module, three genes (*COL3A1, COL1A2,* and *JCHAIN*) have opposite co-expression relationships with the other genes between the normal and tumor conditions (Figure 5C), and these genes are all involved in the regulation of immune response. In particular, since the *COL3A1* and *COL1A2* genes are both in the pathway of scavenging by class A receptors, which are important regulators of immune responses to cancer [74], their co-expression changes in the tumor cells may help us better understand the immune response processes in breast cancer.

### scLink identifies gene network changes from time course data

To further demonstrate scLink’s ability to quantify gene co-expression strength and infer gene networks in single cells under different conditions, we applied scLink to a total of 758 single cells profiled by scRNA-seq (Fluidigm C1 platform) at 0, 12, 24, 36, 72, and 96 h of definitive endoderm (DE) differentiation [75]. We first compared the gene-gene correlation calculated by scLink between 51 lineage-specific marker genes [75] at different time points in the differentiation process (Supplementary Figure S10A). Among the 1275 marker gene pairs, 240 pairs have a correlation change greater than 0.5 along the time course of DE differentiation (Supplementary Figure S10B). For example, the two genes *NANOG* and *PECAM1* are weakly associated at the early time points (0, 12, and 24 h), but moderately associated at the late time points (36 and 72 h) (Figure 6A); the genes *MT1X* and *SOX17* do not demonstrate association at the early time points, but become negatively associated at 36 and 72 h (Figure 6B). These findings are consistent with previous observations in DE differentiation studies [76, 77], but our results provide a detailed view of the association changes between the time points, thanks to the availability of the time course scRNA-seq data. These results may be used to interpret how these genes jointly regulate cell fate decisions in DE differentiation. In contrast, the Pearson correlation coefficients between these two gene pairs remain constantly low at almost all time points (Supplementary Figure S10C), making it difficult to identify and interpret the association changes along the time course.

**Figure 6:**
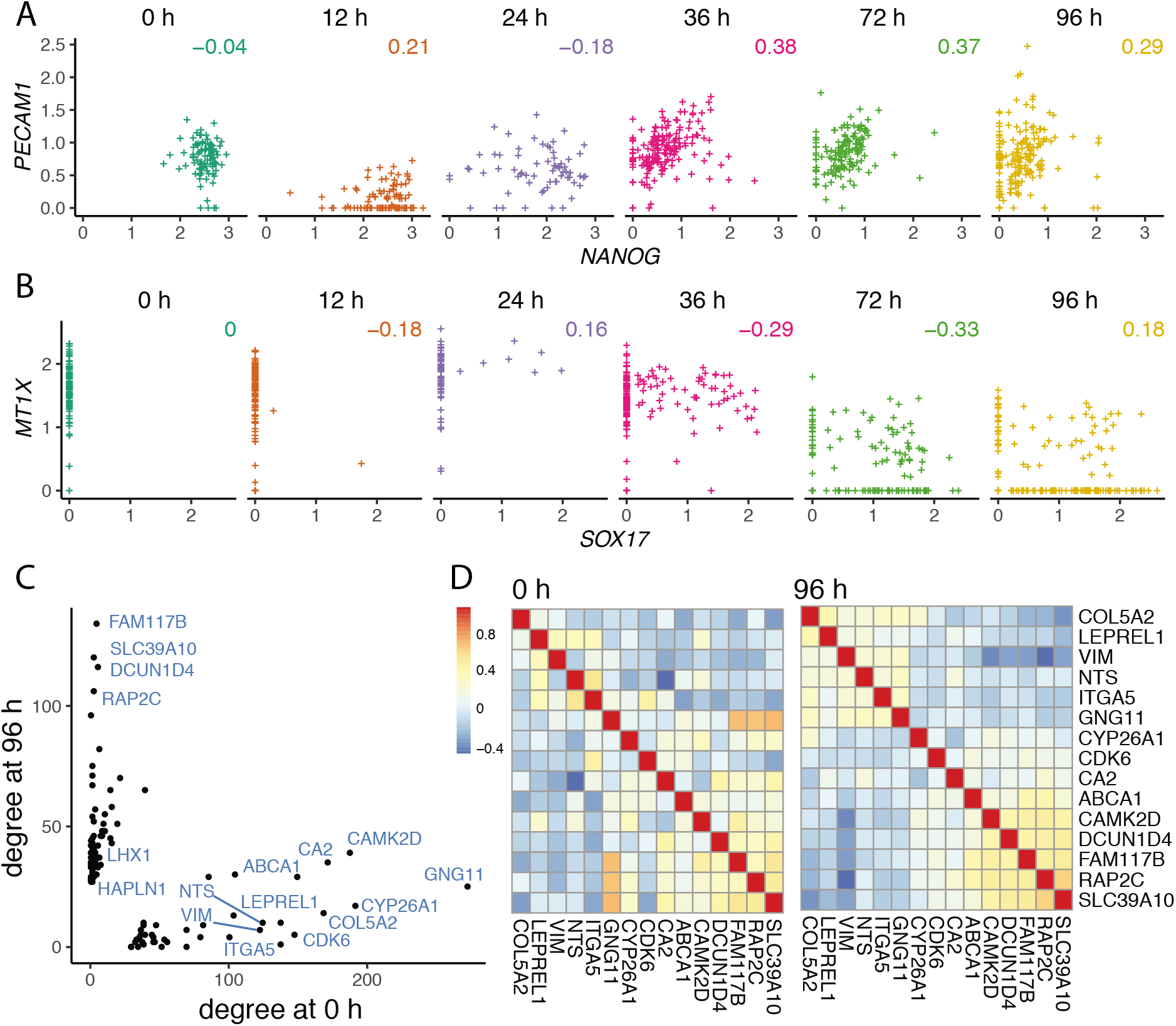
scLink identifies gene network changes along the time course of DE differentiation. **A-B**: The co-expression between *PECAM1* and *NANOG* (**A**), *MT1X* and *SOX17* (**B**) at different time points in the differentiation process of DE. Displayed numbers are correlation measures calculated by scLink. **C**: For genes whose degree changes are greater than 25 between 0 h and 96 h, their degrees at both time points are displayed. Labeled genes have degree changes greater than 100. **D**: scLink’s correlation matrices of genes having degree changes greater than 100 between 0 h and 96 h.

Next, we separately applied scLink to the scRNA-seq data from 0 h and 96 h, using the top 1000 highly expressed genes in the whole dataset. The proportions of zero counts in the two expression matrices are 1.8% and 2.1%, respectively, and the gene-level zero proportion is 0 ~ 98.9% at 0 h and 0 ~ 46.8% at 96 h. The regularization parameters (λ) are selected as the smallest value from {0.2,0.19,…, 0.01} such that the inferred network has no more than 1% edges (i.e., 1% × (1000 × 999)/2 = 4995 edges). In the 0 h gene network, the gene degree ranges from 2 to 274 with an average of 11.9, and the 188 communities identified by the Louvain algorithm has an average size of 5.3. In the 96 h gene network, the gene degree ranges from 2 to 136 with an average of 11.8, and the 296 communities identified by the Louvain algorithm has an average size of 3.4. By comparing the gene networks identified for the two time points, we found 595 differential edges whose corresponding gene pairs have a greater than 0.5 change in their co-expression (Supplementary Figure S11).

To compare the differences in hub genes between the two time points, we assessed the change of gene degrees from 0 h to 96 h. We found that genes with higher degrees at 0 h are enriched with GO terms relevant to mitosis, cell cycle, and chromosome separation, while genes with higher degrees at 96 h are enriched with GO terms relevant to regulation of cell differentiation and organismal development (Supplementary Figure S12). Among the genes that have the largest changes in their degrees, we observed three lineage-specific marker genes, *LHX1, HAPLN1,* and *GNG11* (Figure 6C). In addition, among the genes that have much larger degrees at 0 h than at 96 h, we observed *CYP26A1, CDK6, VIM,* and *ITGA5,* which were shown to have regulatory roles in cell proliferation and/or cell differentiation (Figure 6C) [78, 79, 80]. Visualizing the pairwise coexpression matrices between these genes, we can observe two tight gene modules at 96 h but only one such module at 0 h (Figure 6D), implying that the joint expression of the genes including *COL5A2, LEPREL1, VIM, NTS, ITGA5,* and *GNG11* may be critical for the differentiation of embryonic stem cells. The above results show that the gene co-expression networks identified by scLink from different time points provide important clues regarding transcriptional changes in the differentiation process of DE. In contrast, we also constructed Pearson correlation networks with the same level of sparsity (i.e., 4995 edges), and investigated the biological functions of genes with high degrees at 0 h or 96 h. We found 30.9% of the 1000 genes to have the same direction of degree change in the scLink and correlation networks (Supplementary Figure S13A). Unlike the scLink networks, genes with higher degrees at 0 h in the correlation network are enriched with general GO terms of translation processes, while genes with higher degrees at 96 h are enriched with GO terms relevant to apoptosis (Supplementary Figure S13).

### scLink demonstrates computational efficiency and robustness

To evaluate the computational efficiency and robustness of gene network construction, we compared the performance of scLink with PIDC and GENIE3 based on scRNA-seq data from two technologies, Smart-Seq2 [46] and 10x Genomics [81]. For the Smart-Seq2 technology, we selected four cell types, late pro-B cells, bladder urothelial cells, myeloid cells, and microglial cells, from the Tabula Muris database. The cell numbers of the four cell types are 306, 684, 1208, and 4394, respectively. For each cell type, we selected the top expressed 100, 200, and 500 genes for network construction to evaluate how computational efficiency depends on the network scales. In order to assess the robustness of the three methods given random variation, for each cell type, we randomly selected half of the cells for network construction and independently repeated the procedure ten times with each method. The robustness score of each method was calculated based on the consistency between the ten inferred networks of the same cell type (see Methods). It is a score between 0 and 1, with 0 indicating non-overlap between the ten networks and 1 indicating complete overlap. For each method, the summarized computation time and memory usage were averaged across the ten repeated experiments. Our results show that scLink achieves higher robustness than PIDC and GENIE3 while requiring much less computation time and memory usage (Supplemental Figure S14). For the 10x Genomics technology, we evaluated scLink and PIDC based on scRNA-seq data of 10,085 B cells [81]. Since the computation time of GENIE3 for Smart-Seq2 data exceeded 10^5^ s (2.8 hours) when 500 genes and 2197 cells were used, we did not test it on the large-scale 10x data. For the B cells, we selected the top expressed 500 and 1000 genes for network construction. In order to assess the robustness, we randomly selected 5000 or 8000 B cells for network construction and independently repeated the procedure 10 times for each method. The results again demonstrate that scLink achieves higher computational efficiency and robustness than PIDC (Supplemental Figure S15). In addition, scLink can finish the computation in fewer than 100 s to construct a co-expression network of 1000 genes from 8000 cells. We also notice that the best robustness score achieved by scLink is around 0.5, and this could be explained by two major reasons. First, the correlation calculation and network inference are inevitably affected by the random variation in single-cell gene expression data, leading to variation of identified edges for randomly sampled cells of the same cell type. Second, since the cell types in real scRNA-seq data were also computationally inferred, there may exist cell subtypes that have biologically different gene co-expression networks. The above experiments were performed using the Ubuntu 16.04.5 system and two 8-core CPUs of Intel Xeon CPU E5-2670 at 2.60GHz.

## Discussion

In this work, we developed a method called scLink to improve the construction of sparse gene co-expression networks based on single-cell gene expression data. To demonstrate the applications of scLink and disseminate the research findings in our real data studies, we also developed a web application of scLink (https://rutgersbiostat.shinyapps.io/sciink/). This application provides an interactive platform for users to subset and visualize the cell-type-specific correlation matrices and gene networks constructed by scLink (Supplementary Figure S16). For easy application of scLink to additional single-cell gene expression datasets, we also implemented the methods in the R package `scLink` (https://github.com/Vivianstats/scLink).

In the simulated and real data studies, the gene networks we constructed contain 100 to 1000 genes depending on the dataset used. In actual applications of scLink, we also suggest a filtering step of genes based on the gene detection rates and/or the mean expression levels [16]. Instead of selecting an arbitrary number of genes to be retained, researchers can also set the threshold such that genes of particular interest will be included. The rationale for implementing this filtering step is that genes with small detection rates and low expression level often have low biological relevance and do not provide sufficient information for co-expression estimation, and including these genes might increase false edges in the gene networks. For example, our simulation studies demonstrate that the accuracy of network construction decreases with increasing level of sparsity in single-cell data, regardless of the method being used (Figure 2). Given the relatively high sparsity level of data generated by droplet-based scRNA-seq protocols [82], this filtering step is especially necessary on gene expression datasets from these protocols. When it is of interest to construct a network of thousands of genes, it is still possible to directly apply scLink, but the likelihood optimization step would be more time-consuming because it involves large scale matrix operations. An alternative approach is to first divide the genes into a few major modules based on scLink’s correlation, then apply scLink to each major module separately to identify co-expression networks.

Since the first step of scLink was partially motivated by our scImpute method [29] and additional imputation methods for single-cell gene expression data have also become available [83], an alternative approach to the construction of gene co-expression networks is to apply conventional inference methods on imputed gene expression data. However, we would like to discuss two potential issues with this approach. First, previous studies have shown that imputed data may still be much sparser than bulk data, even though containing fewer zero counts than the observed single-cell data [29, 84]. Therefore, conventional methods designed for bulk data may still have poor performance in network construction even when applied to imputed single-cell data. Second, imputation of gene expression could be a time-consuming step depending on the number of individual cells in the data [85]. By skipping the imputation step and directly accounting for the sparsity issue in co-expression calculation, scLink can achieve better computational efficiency.

Even though our simulation and real data studies demonstrated the great potential of scLink on different types of single-cell gene expression data, we need to interpret the results with caution, since the edges identified by scLink are based on statistical dependencies and do not have directions. These edges may capture the actual regulatory processes, such as the relationships between transcription factors and their putative target genes. However, the inferred edges may also represent co-regulatory relationships of genes regulated by common transcription factors. In addition, we may also identify edges between genes that are responsible for similar biological functions and demonstrate coordinated expression patterns. It is not feasible to directly distinguish the above different types of edges using only gene expression data, but scLink’s results provide good candidates for further computational and/or experimental validations. For example, singlecell ChIP-seq experiments could be designed and prioritized based on scLink’s identified TF-gene pairs [86]. It is also possible to take advantage of existing databases of transcription factors and protein-protein interactions at the validation step, but the knowledge derived from previous bulk tissue research does not necessarily reflect the true scenario in single cells [26].

As discussed in several recent methods, it is possible to more directly infer gene regulatory relationships instead of co-expression relationships on the condition that temporal information is available for the single cells. Some of these methods take pseudo-time orders estimated by other computational methods [87, 27], while others assume actual time course data are available [88]. Since most scRNA-seq experiments were not performed along a time course, only pseudo-time orders may be available for the majority of datasets. However, since pseudo-time orders are only point estimates of physical time orders, it is important to consider how to quantify pseudotime uncertainty and propagate this into the construction regulatory gene networks [89]. Aside from the temporal information, additional experimental data, such as ATAC-seq or ChIP-seq data, have also been shown to assist the inference of gene regulatory networks in bulk tissue studies [90, 91]. As single-cell multi-omics technologies and data integration methods continue to emerge and evolve [92, 83], it will become possible to modify and extend existing bulk-tissue methods for single-cell data. The penalized likelihood approach used in scLink will be able to incorporate the above additional information with flexibility. For instance, we can extend the penalty terms in scLink to apply different levels of regularization on each gene pair based on the epigenetic or chromatin accessibility information. Another future direction for extending scLink is to construct differential gene networks between biological conditions from scRNA-seq data. In our real data studies, we used a straightforward approach to identify differential edges based on the differences in scLink’s correlation strength. However, it is possible to extend the likelihood model to directly identify differential networks using scRNA-seq data from both conditions, as previously done for bulk tissue RNA-seq data [31, 93, 94]. With the ongoing efforts of single cell atlases such as the Human Cell Atlas [95] to better define cell types, states, and lineages, it will also become possible to investigate how gene co-expression and interactions differ in related tissue and cell types. In summary, we expect scLink to be a useful tool for inferring functional gene networks from singlecell gene expression data, with the potential to incorporate other omics data types as single-cell technologies continue to develop.

## Conclusions

In this work, we developed a method called scLink to improve the construction of sparse gene co-expression networks based on single-cell gene expression data. In the scLink method, we first propose a new correlation measure for the strength of gene co-expression relationships, while accounting for the sparsity feature of single-cell gene expression data. Next, relying on the more robust correlation measure, scLink identifies gene co-expression networks in single cells using a penalized and data-adaptive graph model. We first evaluated and compared scLink with five other state-of-the-art methods using carefully designed synthetic networks and gene expression data. These alternative methods include two methods specifically designed for single-cell data and three conventional methods for gene network inference. Our simulation studies showed that scLink has the best accuracy in gene network construction, given different network topologies (hub-based or power-law), gene numbers, and sparsity levels of gene expression. We then conducted a series of real data studies to evaluate the performance of scLink on real single-cell gene expression data. Our results based on the Tabula Muris database show that scLink is able to identify cell-type-specific networks and functional gene modules, and the edges inferred by scLink can potentially capture regulatory relationships between gene pairs. Moreover, our real data studies also demonstrate that scLink can help identify co-expression changes and gene network rewiring between healthy and disease states such as breast cancer. In addition, scLink was also demonstrated to reveal network differences and critical hub genes in time course data, such as those from the differentiation process of definitive endoderm.

## Availability of data and materials

All single-cell gene expression data analysed during this study are included in published articles and their supplementary information files [47, 13, 75]. The source code used in this article is available at https://github.com/Vivianstats/scLink, which is under GNU General Public License v3.0.

## Authors’ contributions

**Wei Vivian Li**: Conceptualization, Methodology, Formal analysis, Software, Writing - Original Draft;

**Yanzeng Li**: Software, Writing- Reviewing and Editing.

## Acknowledgments

The authors thank Dr. Jian Cao at Rutgers Cancer Institute of New Jersey for the helpful discussions. We also thank the editor and the anonymous reviewers for their insightful comments and suggestions. The research reported in this manuscript was supported by the Rutgers School of Public Health Pilot Grant (to WVL) and the New Jersey Alliance for Clinical and Translational Science Mini-methods Grant (to WVL), a component of the National Institute of Health (NIH) under Award Number UL1TR0030117. Computational resources were provided by the Office of Advanced Research Computing (OARC) at Rutgers, The State University of New Jersey, under the National Institutes of Health Grant No. S10OD012346.

